# Conformational sampling and interpolation using language-based protein folding neural networks

**DOI:** 10.1101/2023.12.16.571997

**Authors:** Diego del Alamo, Jeliazko R. Jeliazkov, Daphné Truan, Joel D. Karpiak

## Abstract

Protein language models (PLMs), such ESM2, learn a rich semantic grammar of the protein sequence space. When coupled to protein folding neural networks (e.g., ESMFold), they can facilitate the prediction of tertiary and quaternary protein structures at high accuracy. However, they are limited to modeling protein structures in single states. This manuscript demonstrates that ESMFold can predict alternate conformations of some proteins, including *de novo* designed proteins. Randomly masking the sequence prior to PLM input returned alternate embeddings that ESMFold sometimes mapped to distinct physiologically relevant conformations. From there, inversion of the ESMFold trunk facilitated the generation of high-confidence interconversion paths between the two states. These paths provide a deeper glimpse of how language-based protein folding neural networks derive structural information from high-dimensional sequence representations, while exposing limitations in their general understanding of protein structure and folding.

## 1 Introduction

Dynamics allow proteins to carry out complex biological functions [1, 2, 3, 4], but cannot be reliably predicted from sequence alone [5]. Tuning the inputs of the alignment-based protein folding neural network AlphaFold2 [6] can sometimes accurately model proteins in multiple states [7, 8, 9, 10, 11]. However, these approaches require significant manual intervention and scale poorly to the proteome-level [12, 13, 14], precluding large-scale analyses of protein dynamics akin to those recently carried out on millions of static models [15, 16, 17]. Recently, protein folding neural networks coupled to large PLMs have achieved nearly state-of-the-art performance on structure prediction, particularly on orphan proteins and *de novo* designed proteins, at far faster compute speeds [18, 19, 20, 21]. Yet despite the widespread attention given to emergent qualities of their PLMs [22, 23, 24], their suitability in modeling conformational dynamics remains, to our knowledge, unexplored.

Here we show that the language-based protein folding neural network ESMFold can sample conformational landscapes of some *de novo* designed proteins and natural proteins, pointing to a deeper understanding of protein folding and thermodynamics than previously assumed. Randomly masking amino acids prior to PLM input allows ESM2 to generate alternate residue representations, which ESMFold converted to distinct conformational states. Inversion of the folding trunk allowed interpolation of these representations along high-pLDDT transition paths, thereby generating hypotheses of conformational interconversion (Figure 1). Sequence representations that mapped to distinct conformations of natural proteins, but not *de novo* designed proteins, were highly segregated, with abrupt transitions manifested in high-RMSD structural changes that sometimes skip over high-energy transition states. These results suggest that language-based protein folding neural networks are equipped to rapidly generate hypotheses about protein dynamics. At the same time, they expose discontinuities in how ESMFold maps sequences to representations to structures.

**Figure 1:**
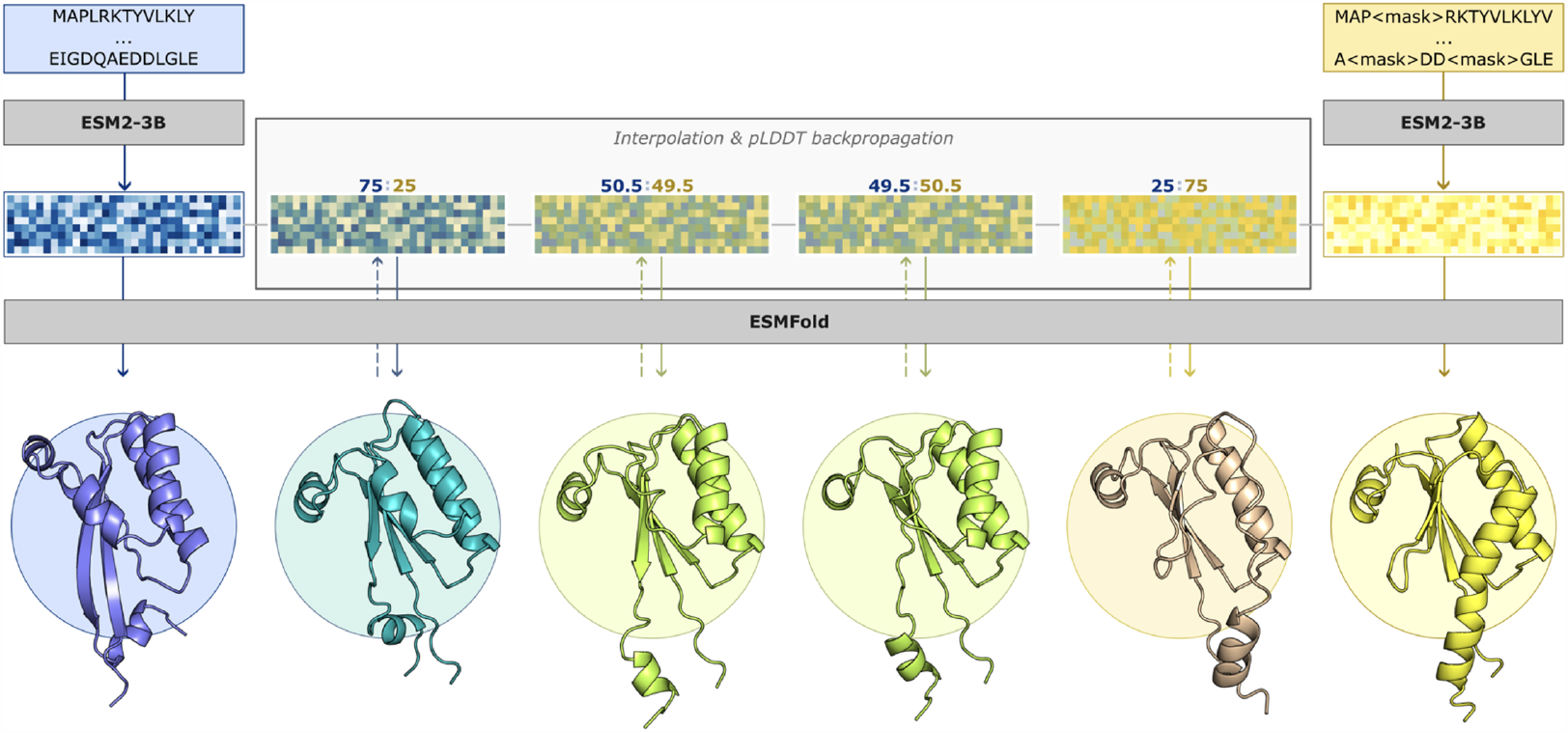
Overview of the conformational sampling and interpolation pipeline. Residue-level representations, shown as heat maps, for initial start and end states are generated using either an unmasked query sequence (left) or a partially masked sequence (right), thereby generating multiple states. Element-wise interpolation of these representations is combined with backpropagation through the ESMFold folding neural network to generate high-confidence transition paths.

## 2 Results and Discussion

### 2.1 *De novo* designed proteins can be modeled in apo and holo conformations

Self-supervised training of large language models, including PLMs like ESM2, relies on randomized masking of tokens in a sequence, with the objective of recovering the unmasked query sequence [25, 26, 27]. Recently, Hermosilla *et al* demonstrated that ESMFold can correctly fold *de novo* designed proteins when large fractions (80-90%) of the query sequence are masked [28]. We reasoned that this approach may be able to sample both states of a recent set of six proteins that were *de novo* designed to interconvert between apo and holo conformations [29].

Both conformations were predicted at high accuracy for two of the six proteins, with the remaining four sampling conformations with features from both states (i.e., putatively intermediate; Figures 2 and S1). For the successful cases, conformational sampling occurred even when only 10% of the sequence was masked. The ESM2 representations of the masked sequences were largely identical to those of the unmasked sequence, with high cosine similarities and low Euclidean distances (Figure S2). Dimensionality reduction of these representations using t-SNE [30] further showed how distinct conformational states were not derived from distinct representations. These results suggested that ESMFold may have learned to extrapolate conformational dynamics in some proteins absent from the training set.

**Figure 2:**
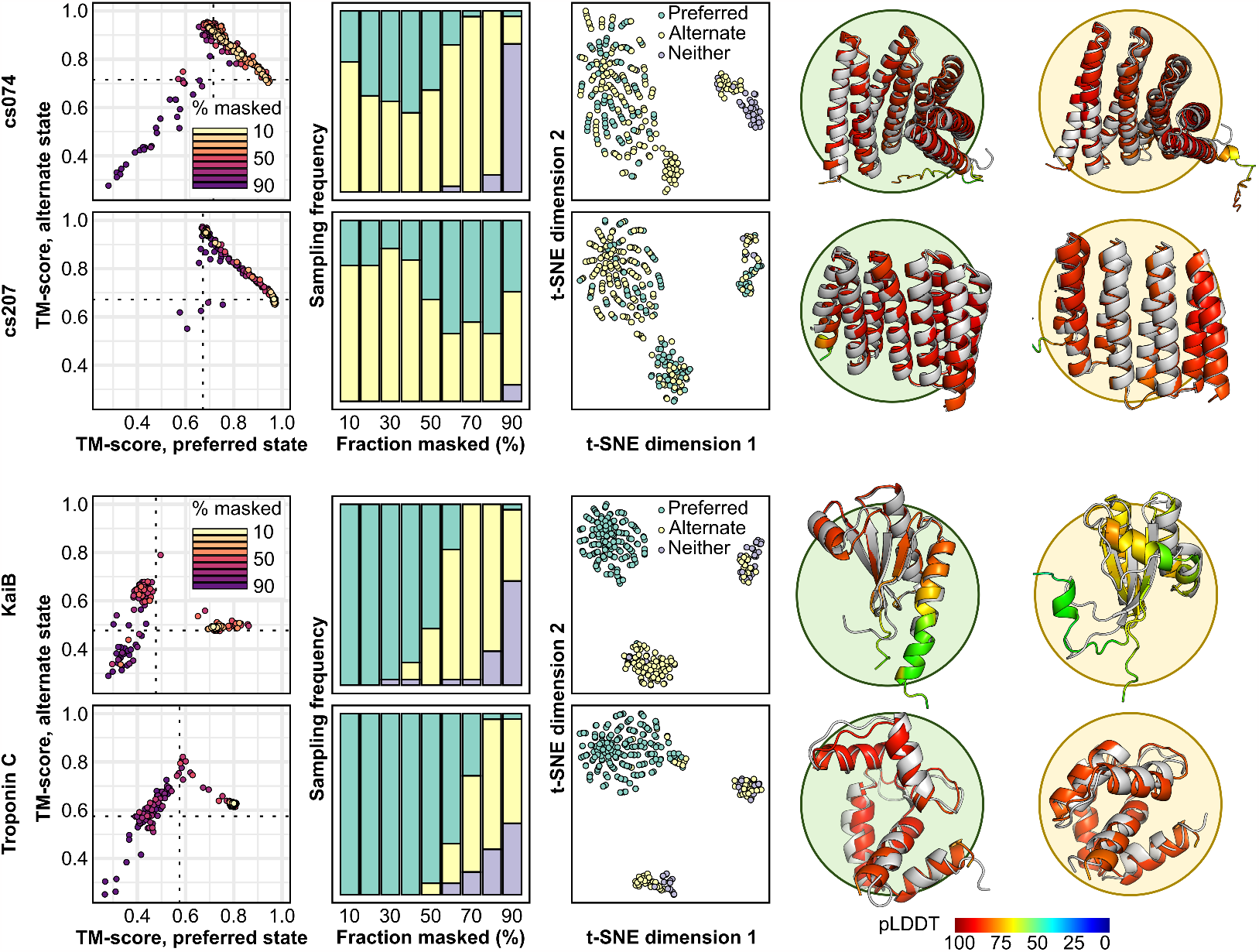
ESMFold predicts alternative conformations of *de novo* designed and some natural proteins. For *de novo* designed proteins (top two rows), no correlation between sequence masking and conformational sampling is observed. In contrast, natural proteins (bottom two rows) only sample alternative states when most of the sequence is masked. Dashed lines indicate TM-scores between reference structures. Cartoon diagrams depict the preferred (left) and alternative (right) states, with reference structures shown in gray. Computational models of the protein cs074 were used as references due to lack of experimental structures.

### 2.2 Distinct conformations of natural proteins are segregated in the embedding space

Expecting these results to be recapitulated in natural monomeric proteins with well-defined dynamics, we assembled a panel of 11 fold-switching proteins, 6 transporters, and 90 ligand-binding proteins from a variety of previous benchmarks [7, 31, 32] (Table S1). Although this approach successfully sampled multiple conformations of some proteins (examples shown in Figure 2), in most cases it failed to do so. Such proteins were predicted either exclusively in one state or, more frequently, in a putatively intermediate state that blended structural features from both states (Figures S3, S4, S5, S6, S7, S8, and S9).

How ESMFold mapped the sequence representations of natural proteins to structure diverged from its treatment of *de novo* designed proteins in several key respects. First, conformational sampling of natural proteins steadily increased with masking rate until a critical threshold appeared to be reached, at which point fewer native-like structures were being generated at all. Second, the ESM2 embeddings for the two states segregated into distinct clusters, with folding failures dispersed elsewhere. Finally, major changes in the per-residue representations were observed in embeddings that yielded the alternative state relative to the unmasked query sequence (Figure S2). The magnitude of these residue-level changes did not appear to correlate with either LDDT [33] (Figure S10) and only weakly correlated with residues’ movement in Cartesian space (Figure S11), consistent with encoding of structural changes at the whole-sequence rather than residue level.

### 2.3 Tracking the determinants of conformational sampling in ESMFold

Why did sequence masking induce conformational transitions in some proteins, but not others? Structural models matching the alternate conformation could be found in the ESMFold Metagenomic Atlas [18] using Foldseek [34] for almost all proteins discussed here, indicating that this effect does not arise from the alternate state being unreachable by the folding trunk (Figure S12). It also failed to correlate with the number of hits to the query sequence fetched from the UniRef50 and UniRef90 databases using Jackhmmer [35, 36] (Figure S13). Moreover, conformational sampling failed to correlate with the query sequence’s pseudo-perplexity, a metric capturing how well it is understood by the PLM [18] (Figure S14). These negative results, combined with those observed in four out of six *de novo* designed proteins, suggest that variations in conformational sampling may arise from the cumulative effects of idiosyncrasies and imbalances in the training set, rather than specific properties of the individual proteins being predicted.

### 2.4 Sampling conformational transition paths by interpolating between representations for different states

Because the representations for both end states are identical in shape, we reasoned that transition paths between these states could be generated by iteratively interpolating between their representations and passing the resulting outputs through the ESMFold folding trunk as shown in Figure 1. The full algorithm is detailed in Section 4; briefly, it proceeded recursively, with initial guesses generated by averaging the representations of high-RMSD pairs of consecutive structural models in the transition path and refining by backpropagation through an inverted ESMFold network, an approach inspired by similar methods for protein design [37, 38, 39, 40, 41]. Loss functions included pLDDT as well as Euclidean distance restraints ensuring that the ESM2 representations for new models in the transition path were equidistant from those of the two flanking models used to generate them.

Nine such trajectories were generated in the fold-switching protein KaiB and the calcium-binding protein Troponin C (Figure 3A). All eighteen trajectories limited conformational interconversion to a very minor fraction of the overall Euclidean space separating the two sequence representations, and no correlation was observed between Euclidean distance traversed in the embedding space and RMSD changes in resulting structures (Figure 3B). Most trajectories were found to skip over high-energy conformational transitions entirely in both proteins, even though the traversed distance in Euclidean terms was extremely small (Figure 3C).

**Figure 3:**
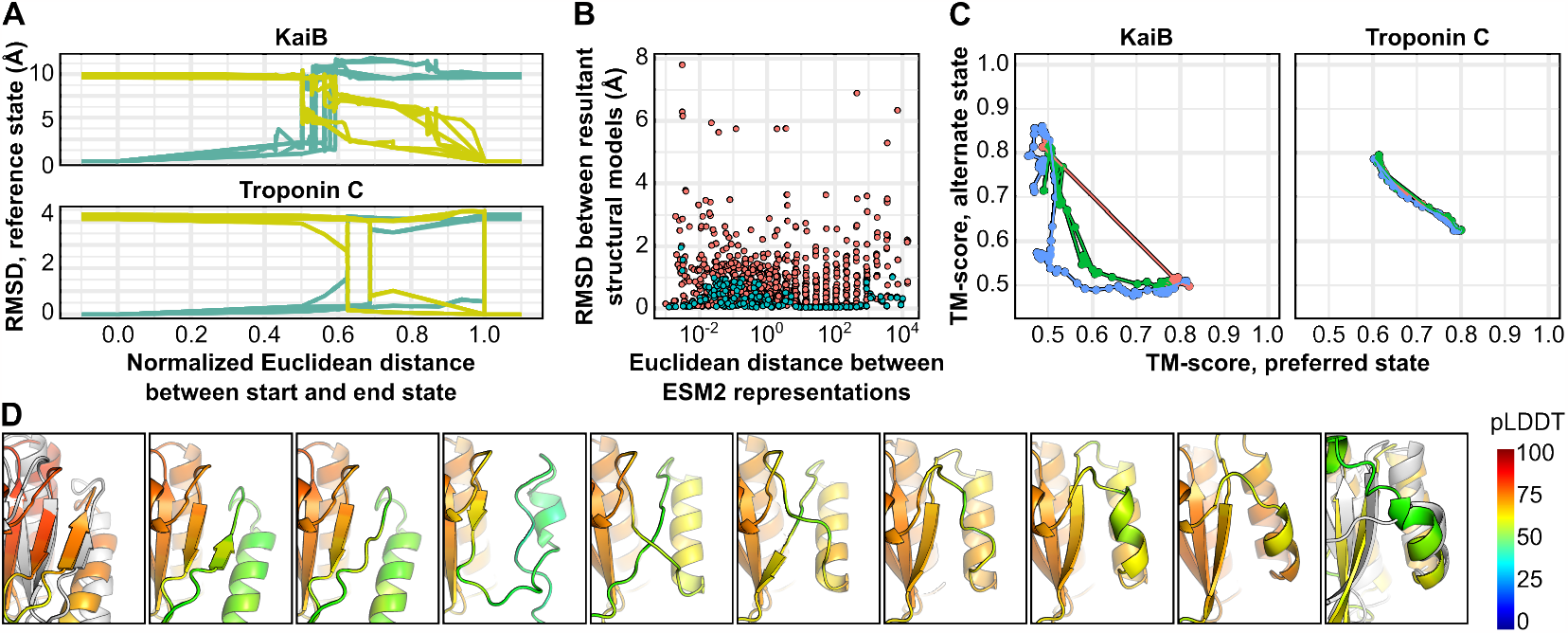
Conformational interpolation between the ESM2 representations of start and end states of KaiB and Troponin C using ESMFold. **(A)** Conformational transitions occur during very short intervals during the transition from one state to another. Start and end points were chosen based on structural similarity to either the preferred state or alternate state and are shown in either teal or yellow. **(B)** Changes in protein representations are not correlated with changes in RMSD in either KaiB (orange) or Troponin C (teal). **(C)** Example trajectories show how different sampling granularities are achieved, with some trajectories skipping over conformational transitions. **(D)** Example sheet-to-helix transition in KaiB residues 62-68, with residues colored by pLDDT. Crystal structures are shown in white on the leftmost and rightmost panels.

Residues 62-68 of KaiB provide an illustrative example, as they undergo a sheet-to-helix transition that must, in principle, sample an intermediate loop conformation. Only one trajectory, shown in Figure 3D successfully sampled such a state. The remaining eight skipped over the state entirely, suggesting that high-energy states occupy an extremely limited fraction of the total embedding space separating the two conformations, and that the boundaries between the representations of the two states do not uniformly reflect biophysically relevant conformational transitions.

## 3 Conclusion

This manuscript demonstrates that language-based protein folding neural networks are primed to explore protein conformational landscapes at scale. Of particular interest was the observation that ESMFold interpreted nearly-identical language model representations of *de novo* designed proteins as distinct conformations, which may point to an emergent understanding of structural dynamics as a fundamental property of some amino acid sequences. However, although we observed success with *de novo* designed proteins that is unachievable with AlphaFold without pipeline modifications [29], the relatively low success rate in our benchmark set proves that reliable conformational sampling is not currently achievable by ESMFold, and may require alternate training schemes to fully unlock [42]. Moreover, our proof-of-concept involving the simulation of transition paths in KaiB and Troponin C demonstrated that the language model’s embedding space does not neatly map to conformational space, as indicated by unphysical transition paths between discrete conformations. Nevertheless, it provides one example of the broader range of hypotheses that could be generated by a more robust language-based protein folding neural network. Finally, while these results are limited to ESM2 and ESMFold, its two-stage training procedure closely aligns with that of other language-based protein folding models such as OmegaFold and RGN2 [20, 43], and similar results would be expected in those models as well.

## 4 Methods

### 4.1 Sequence masking and conformational sampling

Conformational sampling was achieved by modifying a recently described sequence masking approach described in [28]. The identities of a fraction of residues 0.1, 0.2, 0.9 are randomly masked prior to processing into embeddings by the ESM2 language model; our modification is limited to returning the full (*N*_*layer*_, *N*_*res*_, *N*_*dim*_) embeddings for each sequence, rather than the embeddings for the last layer. We applied this to the benchmark set of protein structure pairs in Table S1, which was derived from three benchmarks for conformational change modeling using AlphaFold2 and includes recent *de novo* designed proteins that adopt multiple conformations. Recycles were set to zero.

### 4.2 Interpolation between models in embedding space

Transition paths were generated using a recursive algorithm that interpolates between two sets of language model embeddings *x*_1_, *x*_2_, each of which has the shape (*N*_*layer*_, *N*_*res*_, *N*_*dim*_) and maps to structurally distinct conformers. For the ESM2 model, these representations are of shape 37 * *N*_*res*_ * 2560, or approximately 79, 000 dimensions per amino acid. At each iteration *i*, we seek to introduce a new representative model *x*_*i*_ that interpolates between the highest-RMSD pair of adjacent structures. After calculating pairwise RMSD values for all such pairs along the trajectory, a new representative model is generated by taking the midpoint between the representations for the highest-RMSD pair of consecutive structural models. The resulting calculated representations are passed through the folding trunk of ESMFold to generate a structural model, which is then optimized using two losses: a pLDDT loss equal to negative pLDDT, and a geometric loss *L*_*dist*_ that ensures the new representations are equidistant from the two seed representations:

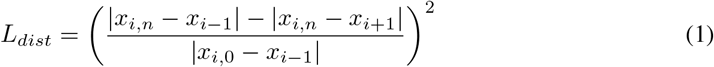

Here, *x*_*i*−1_ and *x*_*i*+1_ are the flanking representations used for interpolation, and *x*_*i*,0_ and *x*_*i,n*_ are the initial and refined embeddings for the model whose position is being interpolated. The Adam optimizer with beta1 = 0.9, beta2 = 0.999 was used with gradient clipping set to 1, and the learning rate was initially set to 1e-3 but was scaled with the distance between the two seed embeddings, *x*_*i*−1_ and *x*_*i*+1_ [44]. Each iteration proceeded until convergence. The algorithm terminated when repeated interpolation failed to decrease the distance between the highest-RMSD pairs of structures. All computations were carried out on an A6000 GPU.

### 4.3 Miscellaneous

Searches using Foldseek (v. 2-8bd520) [34] and Jackhmmer (v3.2.1) [35] were carried out using default settings. t-SNE was used as implemented in python using SciKit-Learn [45]. TM-scores and structural alignments were calculated using TM-align [46].

## Acknowledgments and Disclosure of Funding

The authors would like to thank the GSK Fellows Program for research support and the ESM team at Meta for permissive licensing and development of ESMFold.

## 5 Supplementary Tables

**Table S1:**
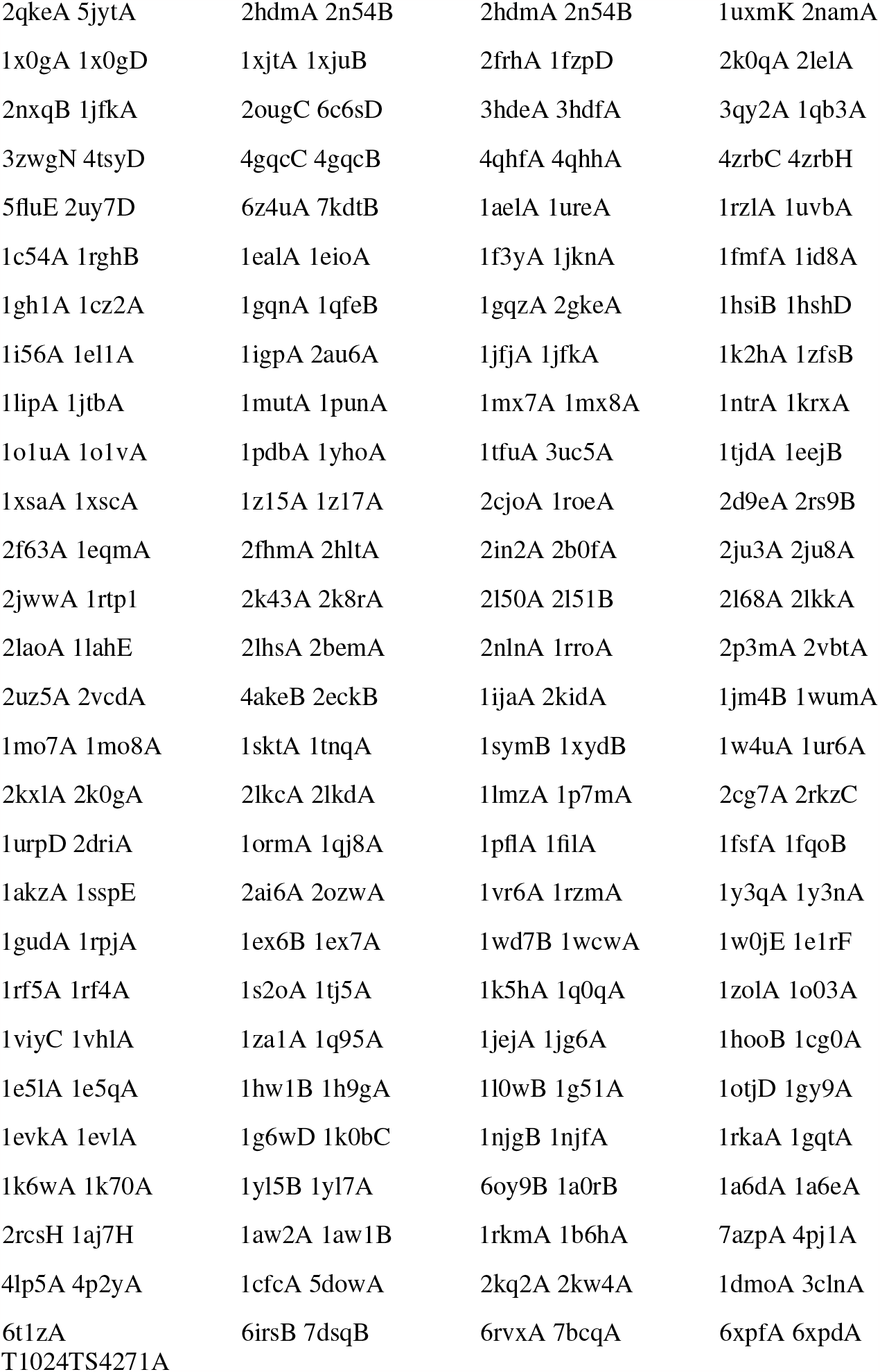
List of PDB pairs used in this study. One of the reference structures for transporter LmrP was derived from CASP14 (T1024TS4271A).

## 6 Supplementary Figures

**Figure S1:**
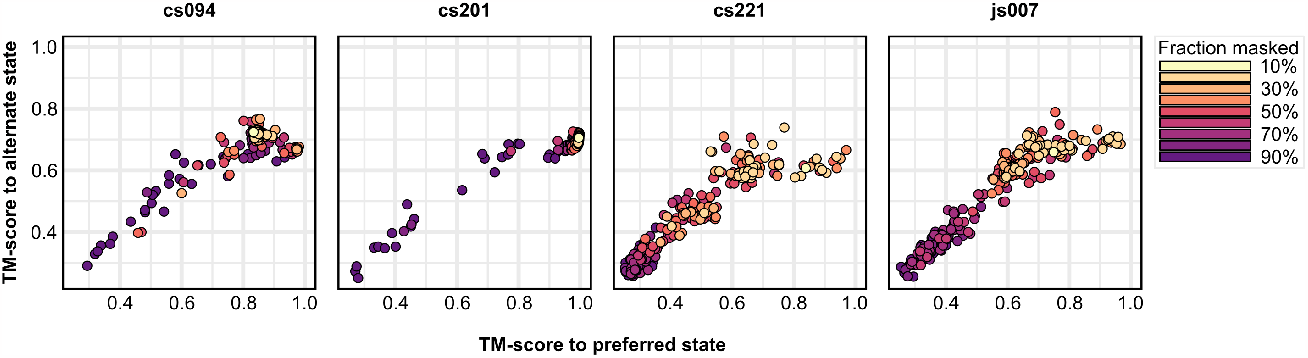
ESMFold predicts intermediate states of some *de novo* designed proteins. Computational models were used for all proteins.

**Figure S2:**
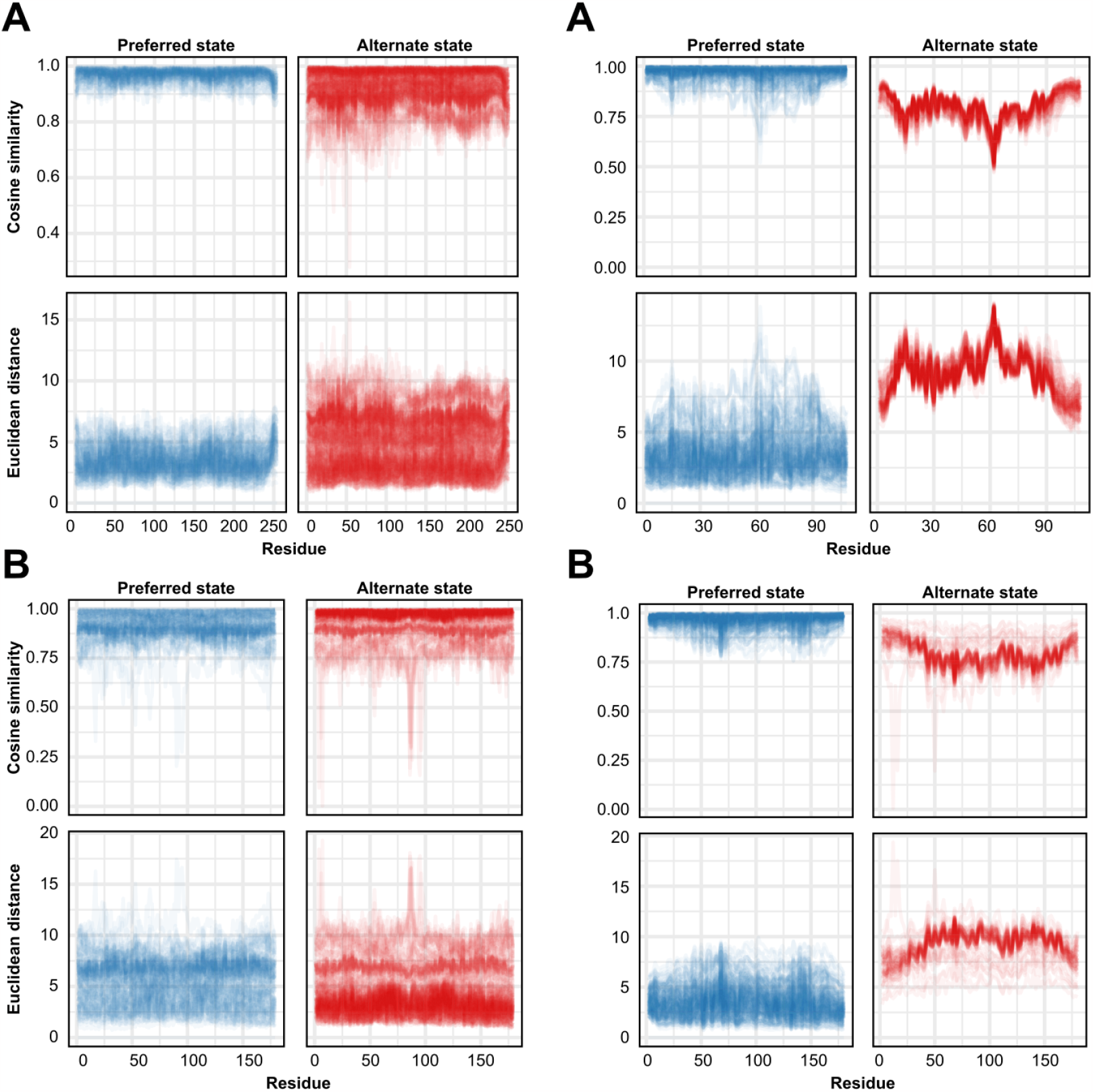
Distinctions in residue-level language model embeddings of designed and natural proteins. The plots show the difference between the unmasked query sequence and those mapping to either the preferred or alternate states of *de novo* designed proteins cs074 (top left) and cs207 (bottom left), and natural proteins KaiB (top right) and Troponin C (bottom right). The representations for alternate conformations of natural proteins on the right show greater dissimilarity from those of the unmasked query sequence than the *de novo* designed proteins.

**Figure S3:**
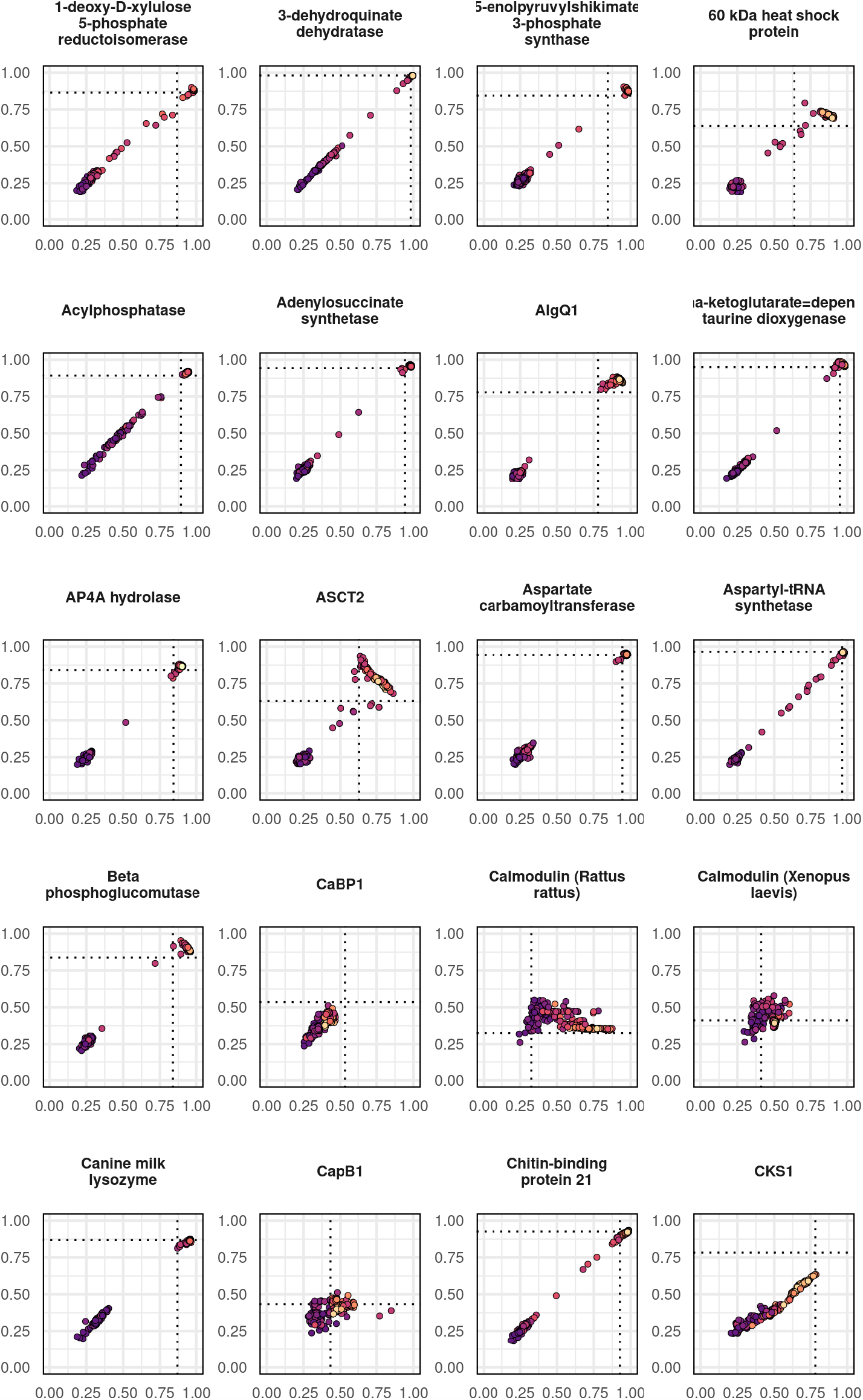
Conformational sampling in benchmark proteins. Preferred and alternate conformations shown on the X- and Y-axes, respectively. Dashed lines indicate TM-scores between reference structures, with a TM-score of 1 indicating that the structures are identical. Colors correspond to the fraction of the amino acid sequence that was masked.

**Figure S4:**
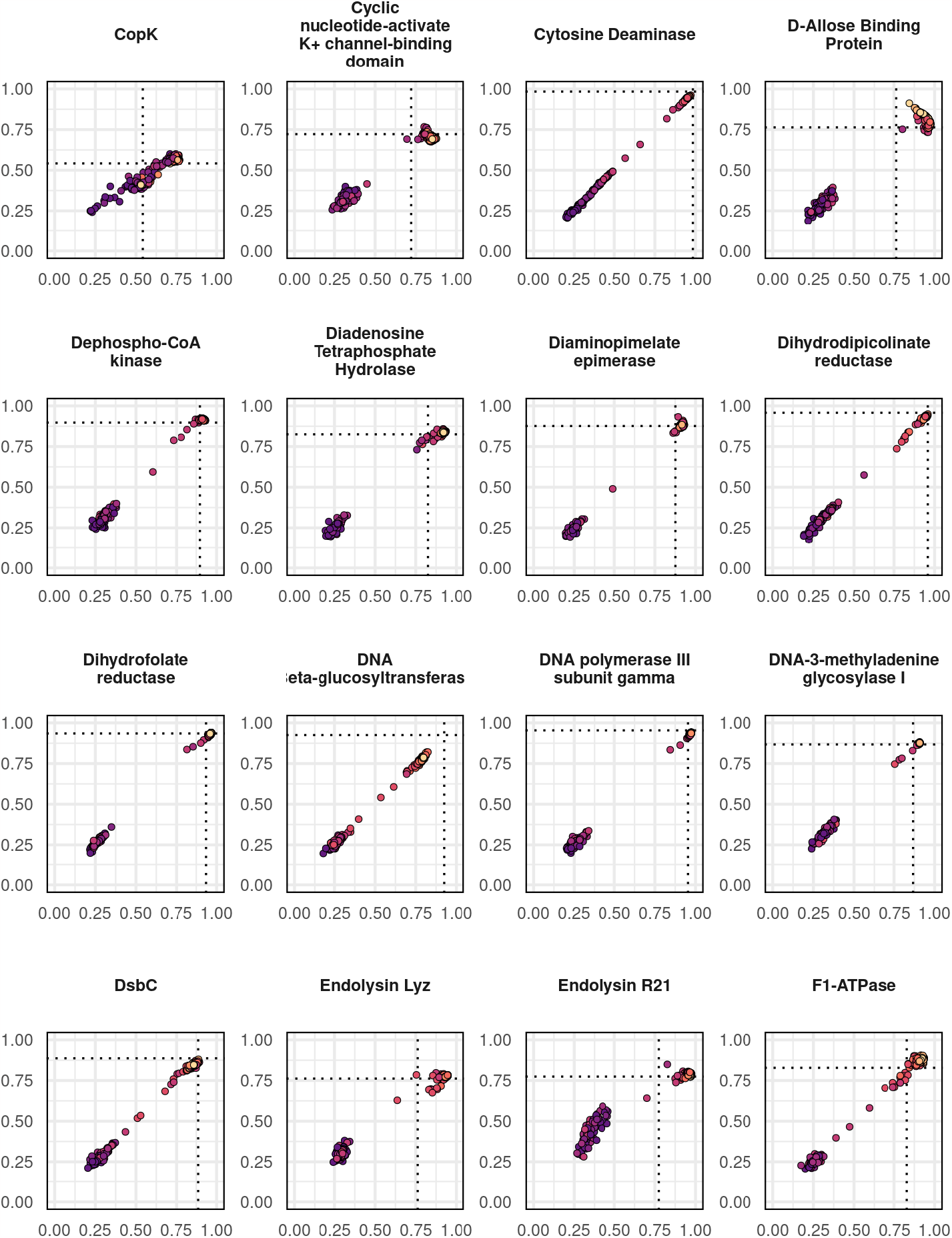
Conformational sampling in benchmark proteins. Preferred and alternate conformations shown on the X- and Y-axes, respectively. Dashed lines indicate TM-scores between reference structures, with a TM-score of 1 indicating that the structures are identical. Colors correspond to the fraction of the amino acid sequence that was masked.

**Figure S5:**
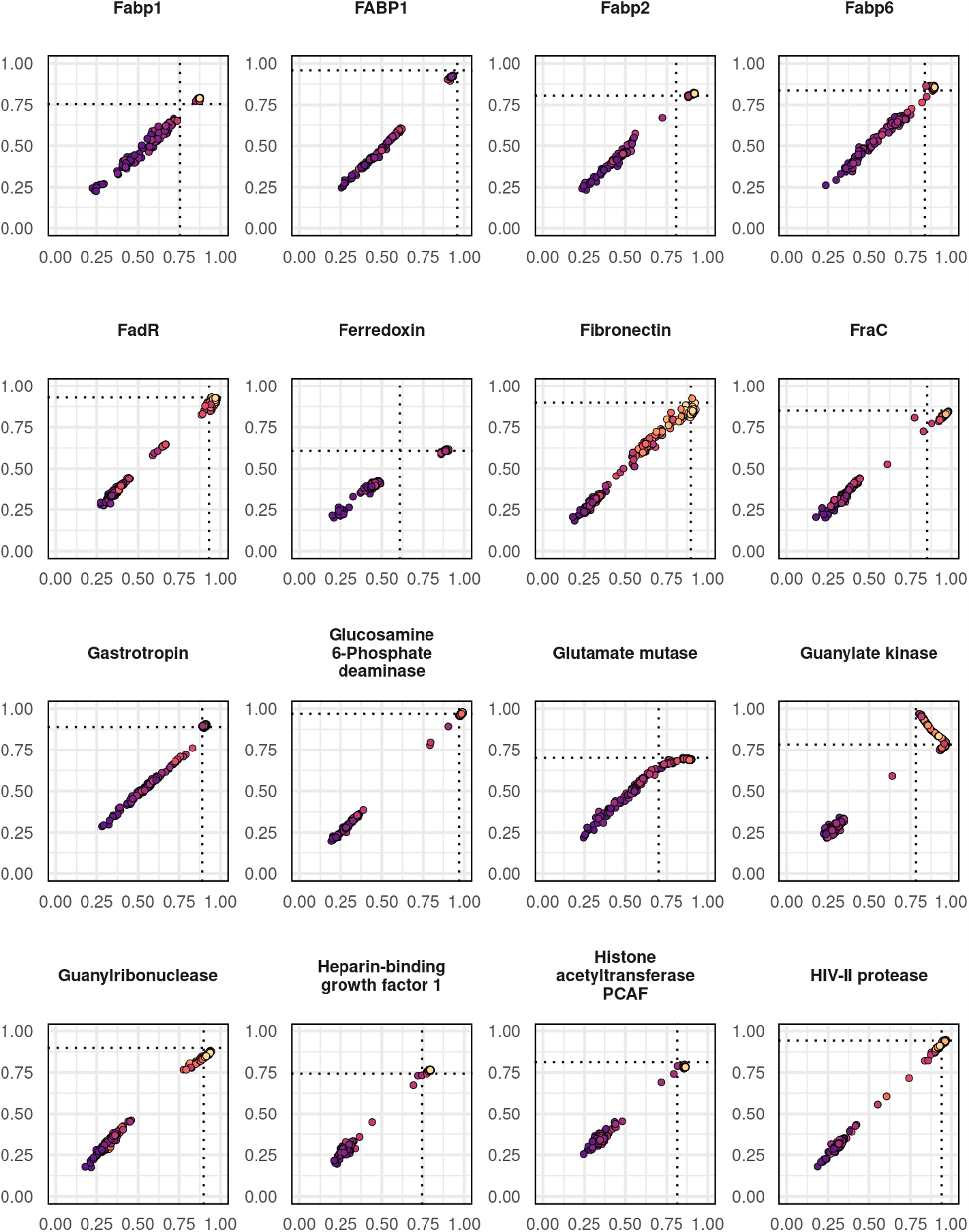
Conformational sampling in benchmark proteins. Preferred and alternate conformations shown on the X- and Y-axes, respectively. Dashed lines indicate TM-scores between reference structures, with a TM-score of 1 indicating that the structures are identical. Colors correspond to the fraction of the amino acid sequence that was masked.

**Figure S6:**
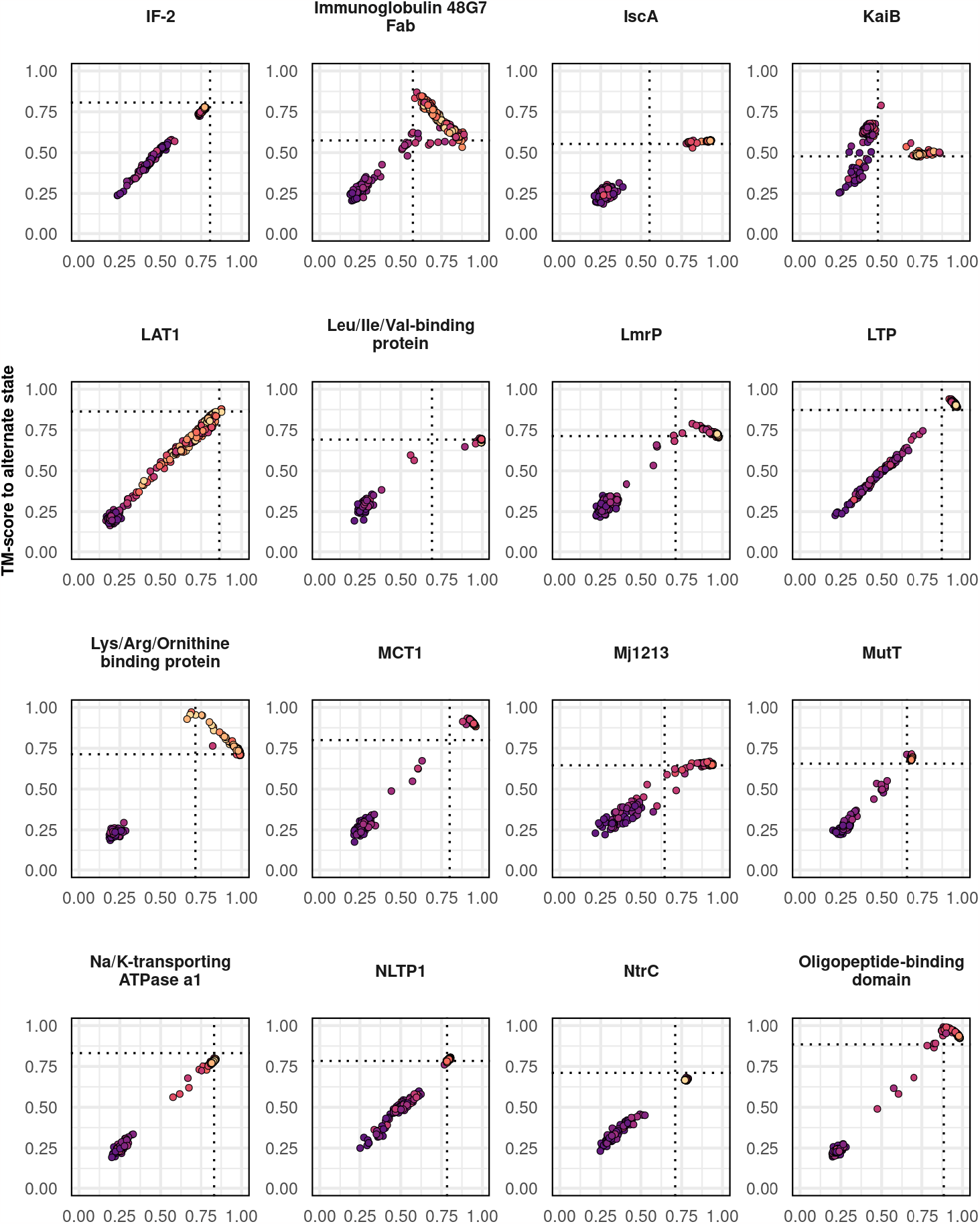
Conformational sampling in benchmark proteins. Preferred and alternate conformations shown on the X- and Y-axes, respectively. Dashed lines indicate TM-scores between reference structures, with a TM-score of 1 indicating that the structures are identical. Colors correspond to the fraction of the amino acid sequence that was masked.

**Figure S7:**
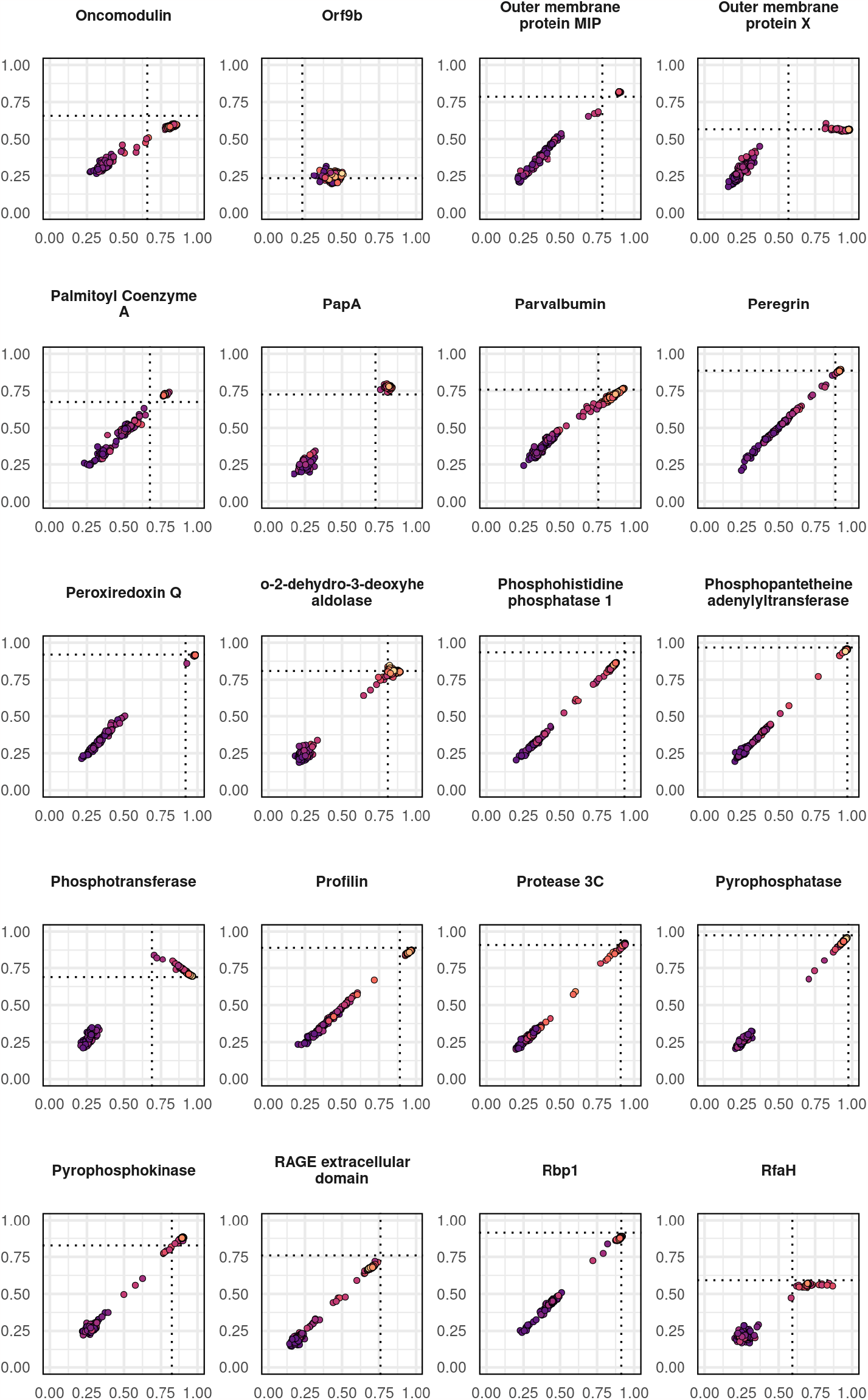
Conformational sampling in benchmark proteins. Preferred and alternate conformations shown on the X- and Y-axes, respectively. Dashed lines indicate TM-scores between reference structures, with a TM-score of 1 indicating that the structures are identical. Colors correspond to the fraction of the amino acid sequence that was masked.

**Figure S8:**
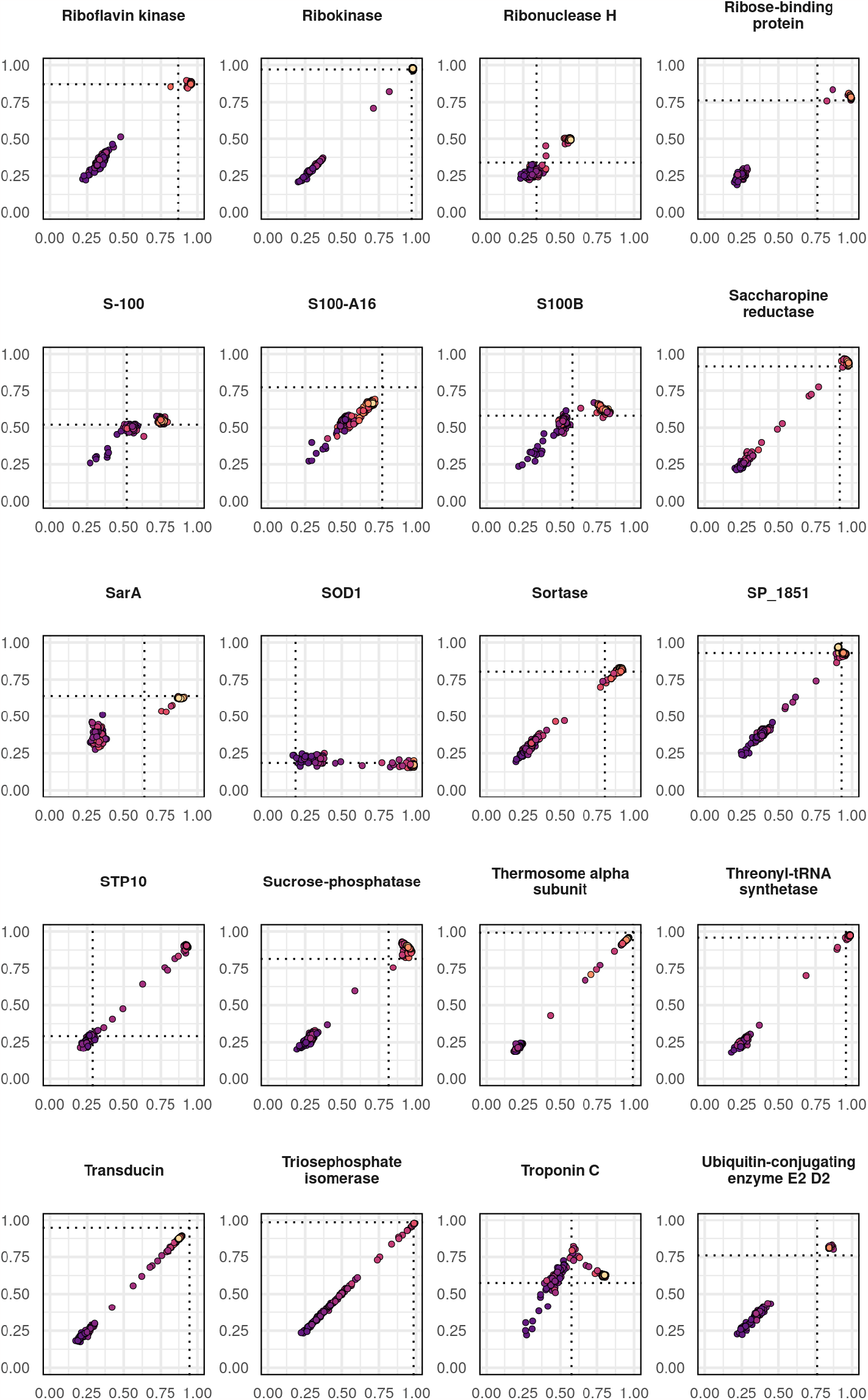
Conformational sampling in benchmark proteins. Preferred and alternate conformations shown on the X- and Y-axes, respectively. Dashed lines indicate TM-scores between reference structures, with a TM-score of 1 indicating that the structures are identical. Colors correspond to the fraction of the amino acid sequence that was masked.

**Figure S9:**
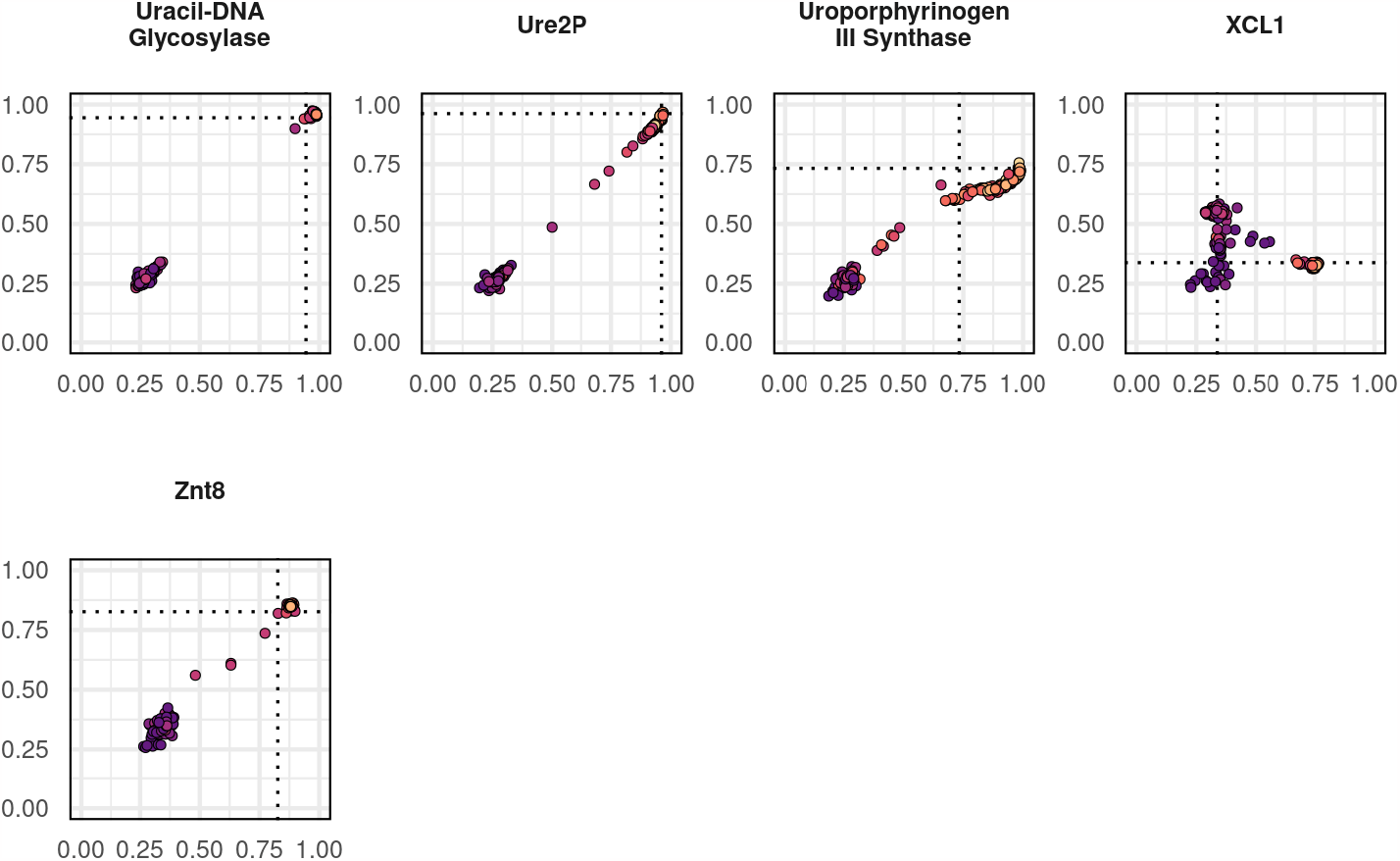
Conformational sampling in benchmark proteins. Preferred and alternate conformations shown on the X- and Y-axes, respectively. Dashed lines indicate TM-scores between reference structures, with a TM-score of 1 indicating that the structures are identical. Colors correspond to the fraction of the amino acid sequence that was masked.

**Figure S10:**
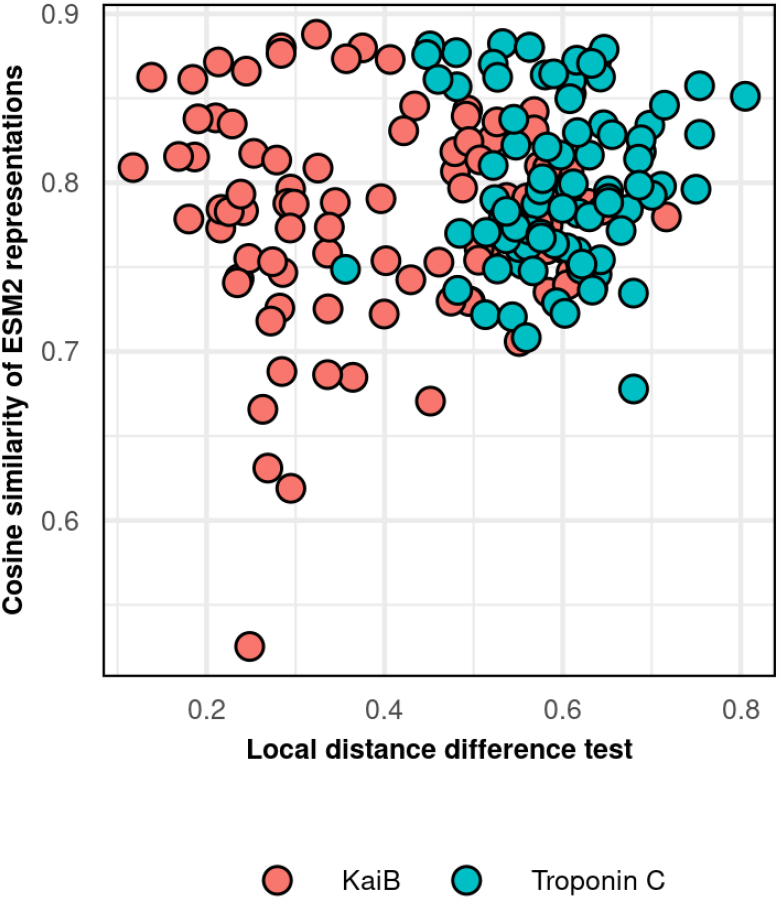
Residue-level structural changes (here, local distance difference test, or LDDT) do not correlate with residue-level differences in the ESM2 representations encoding those conformations.

**Figure S11:**
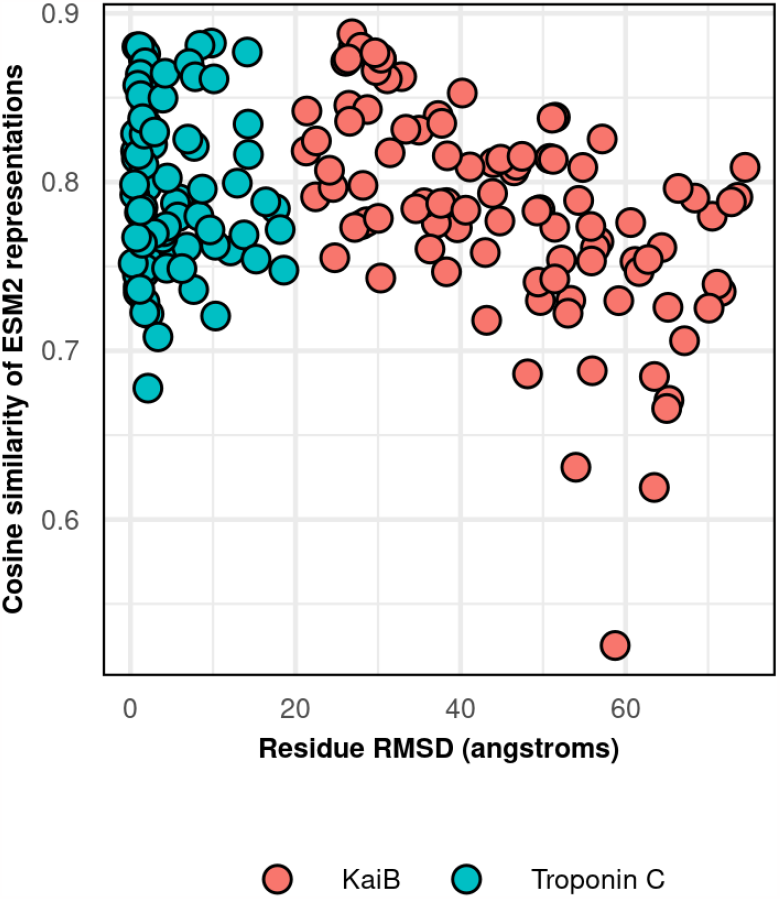
Residue-level structural changes (here, root mean squared deviation, or RMSD) weakly correlate with residue-level differences in the ESM2 representations encoding those conformations in KaiB and not in Troponin C.

**Figure S12:**
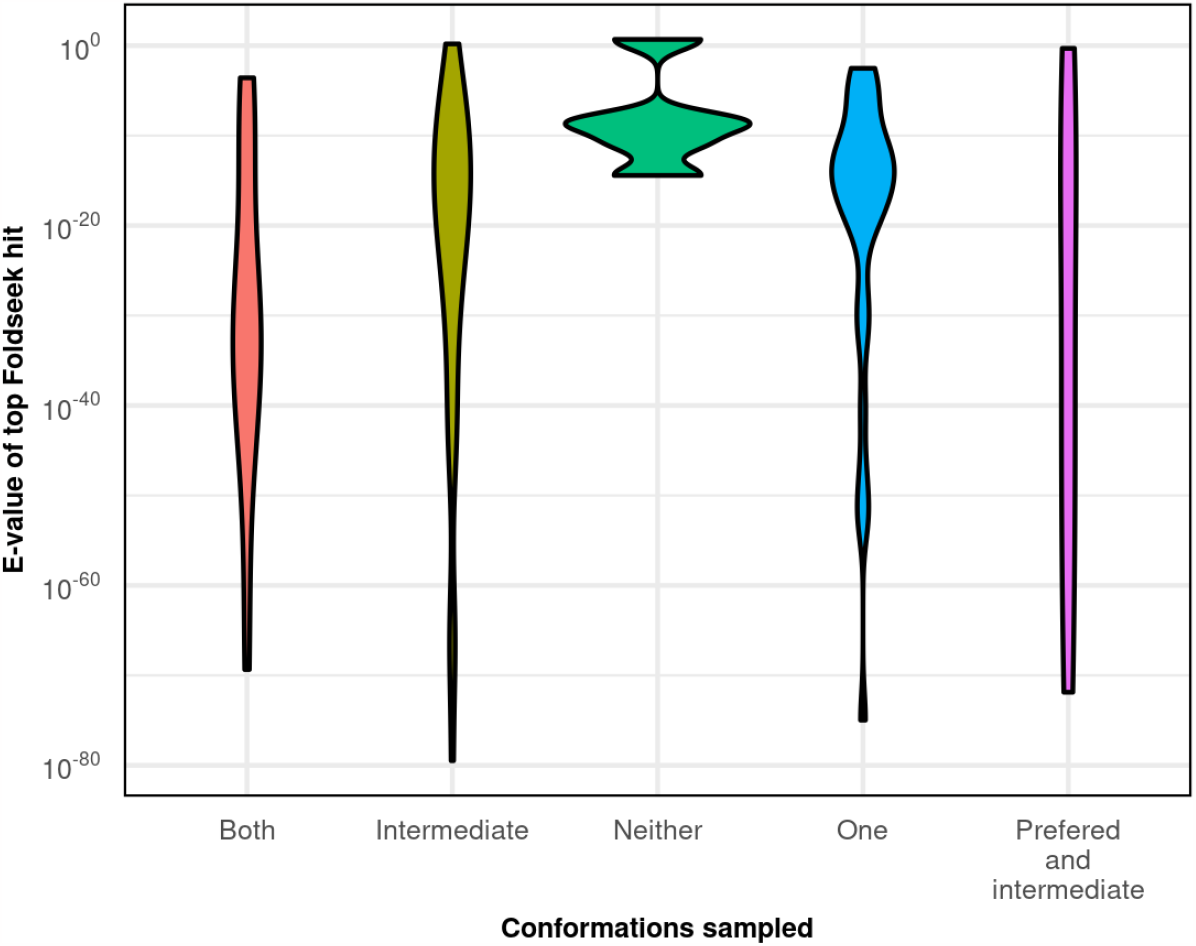
No correlation was observed between conformational sampling success in natural proteins and the presence or absence of similar protein structural models in the ESM Metagenomic Atlas.

**Figure S13:**
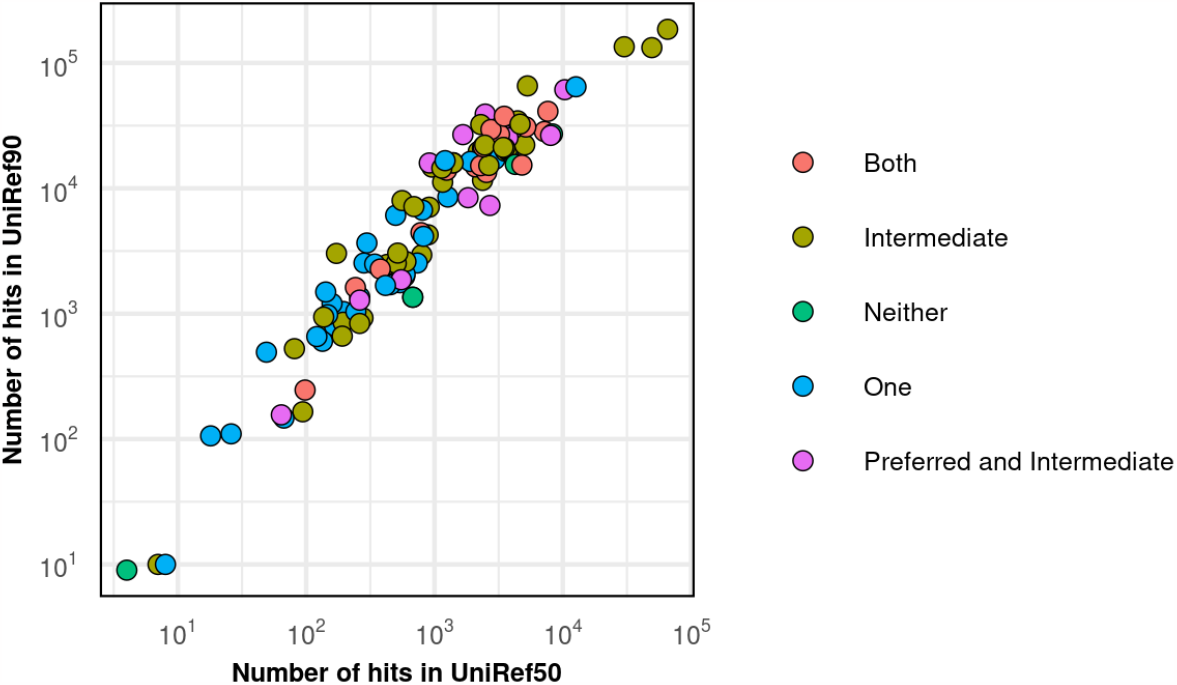
No correlation was observed between conformational sampling using language model masking and the number of hits in either the UniRef50 or UniRef90 sequence databases.

**Figure S14:**
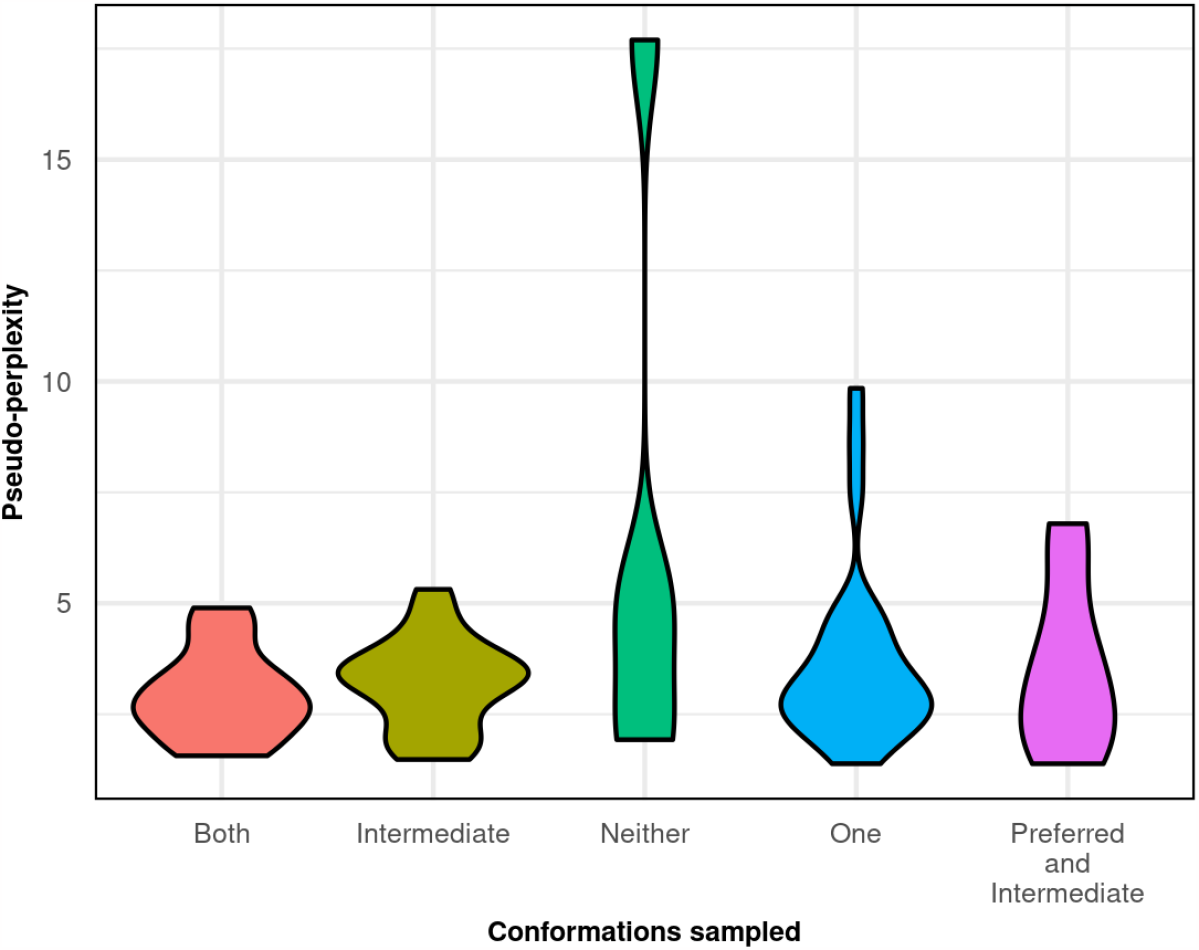
No correlation was observed between conformational sampling using language model masking and the pseudo-perplexity of the query sequences.

